# “Spilling-in” versus “Filling-in”: disentangling bottom-up drive from predictive completion in human V1–V3

**DOI:** 10.64898/2026.10.01.755919

**Authors:** Kelly Chang, Geoffrey M. Boynton, Ione Fine

## Abstract

It is well established that responses in visual areas are a combination of bottom-up stimulus driven input and ‘top-down’ input driven by expectation. However, it has been difficult to quantify the differential strength of these components, and how they vary across the visual hierarchy. Here, we introduce a method for dissociating ‘bottom-up’ and ‘top-down’ contributions in early retinotopic areas (V1-V3). Using a log-scaled bar stimulus, we measured BOLD responses under full-field vs. central scotoma conditions. Using the full-field stimulus, we estimated pRFs in human visual cortex (V1–V3). We then used each voxel’s pRF to predict the response expected from bottom-up responses alone, and compared this prediction to measured responses. Responses within the scotoma demonstrated measurable predictive filling-in that increased systematically from V1 to V3. Our results demonstrate that activity within retinotopic visual cortex cannot be explained solely by feedforward processing and provide a principled method for measuring the contributions of predictive signals in early visual cortex. Finally, we show that top-down filling-in can masquerade as cortical reorganization in which filling-in signals are accounted for through pRF displacements and size increases. Our results show that estimating top-down predictive components is critical for accurately describing cortical responses near regions of visual loss.

## Introduction

When portions of the visual field are absent or occluded, observers often experience perceptual completion of the missing region, a phenomenon broadly referred to as filling-in. Classic examples in neurotypical vision include completion across the physiological blind spot and within artificially induced scotomas, wherein stabilized or masked regions are perceptually replaced by surrounding structure or texture (Komatsu, 2006; Ramachandran & Gregory, 1991; Spillman & De Weerd, 2003; Spillmann et al., 2006).

These neural responses within fMRI population receptive fields (pRFs) that do not receive bottom-up input have three possible causes: (1) they may arise from stimulus-driven responses within pRFs that overlap with the scotoma border – ‘spilling in’, (2) they may reflect top-down filling-in, or (3) they may reflect shifts or expansion of pRF locations to represent regions outside the scotoma.

A large number of human and animal studies have demonstrated top-down filling-in of missing information based on the spatial and statistical structure of the surrounding input (e.g. Muckli et al., 2015; Papale et al., 2023; Smith & Muckli, 2010; Weerd et al., 1995). Predictive coding theories formalize this by proposing that perception arises from hierarchical inference (Friston, 2005; Rao & Ballard, 1999; Roelfsema & de Lange, 2016); higher visual areas generate predictions about sensory input and propagate them to earlier areas via feedback connections (Keller et al., 2020; Kok et al., 2012).

Both theory and the data suggest that the influence of top-down signals should increase across the visual hierarchy. FMRI responses consistent with filling-in have been reported in V1 (Meng et al., 2005; Tong & Engel, 2001), but these signals are relatively weak and are strongly dependent on stimulus context and task demands (Masuda et al., 2008). In contrast, studies of surface completion and illusory contours indicate stronger and more sustained responses in extrastriate areas such as V2 and V3, with context-dependent V1 responses often emerging later in time, consistent with feedback from higher visual areas (Mendola et al., 1999; Weerd et al., 1995).

However, although it has generally been assumed that feedback increases across the visual hierarchy, the relative strength of bottom-up vs. top-down signals across the hierarchy has not yet been quantified. One reason for this is that estimating the relative contributions of bottom-up from top-down signals requires factoring out the impact of differences in receptive field sizes across visual areas. “Filling-in” as a result of top-down signals must be separated from “spilling-in” – the increased spread of neural activity across the cortical surface as a result of larger receptive field sizes.

The second potential source of neural signals within deprived regions of cortex is cortical reorganization – shifts or expansions of receptive fields. Early studies in animals suggested rapid dramatic shifts in V1 receptive fields near the lesion projection zone (Gilbert & Wiesel, 1992; Kaas et al., 1990), although more recent findings that correctly included the presence of the scotoma in the input model found little or no reorganization in V1 (Smirnakis et al., 2005). In humans, pRF shifts and enlargements have been reported near the lesion projection zone in macular degeneration, retinitis pigmentosa and V1 lesions (Baseler et al., 2011; Ferreira et al., 2017; Papanikolaou et al., 2015). However, comparable pRF shifts occur in healthy observers exposed to an artificial scotoma within a single scanning session (Baseler et al., 2011; Binda et al., 2013; Carvalho et al., 2021; Haak et al., 2012; Hummer et al., 2018) or near the rod scotoma under scotopic viewing (Barton & Brewer, 2015). What explains these rapid changes, given that they are unlikely to represent cortical reorganization (Dumoulin & Knapen, 2018; Prabhakaran et al., 2020)? In some cases, these shifts in pRF locations and sizes reflect a failure to include the scotoma in the bottom-up model (Barton & Brewer, 2015; Baseler et al., 2011; Gilbert & Wiesel, 1992; Kaas et al., 1990) as discussed previously (Binda et al., 2013; Papanikolaou et al., 2015). In others it seems likely that top-down predictive signals are mis-modelled as shifts in pRF locations – Binda et al. found shifts near the edge of a scotoma for a predictable stimulus, but not when using an unpredictable multifocal stimulus (Binda et al., 2013). Finally, shifts in in pRFs can occur without changes in the underlying neural receptive fields due to a re-weighting of how the population of neurons within a voxel contributes to the pRF, for example a shift in spatial attention towards the edge of a scotoma might result in neurons on the outer border of the scotoma contributing more heavily to a given pRF response (Dumoulin & Knapen, 2018).

Here we use a combination of retinotopic mapping and forward-modeling to examine responses in an artificial foveal 2-degree scotoma. Having estimated pRFs within V1-V3 we used each voxel’s estimated pRF to predict the bottom-up response – a measure of what proportion of response within the scotoma can be attributed to bottom-up pRF overlap alone - “spilling-in”. The difference between this predicted response and measured signals can be attributed to top-down “filling-in”. We found that the proportion of signal that could be attributed to “filling-in” increased markedly across the visual hierarchy.

We went on to examine the relationship between predictive filling-in and reported shifts in pRF location and size near scotomas. A variety of studies have reported such shifts (Binda et al., 2013; Carvalho et al., 2021; Hummer et al., 2018) but these estimates come from forward models that assume zero input signal within the scotoma. We fit our data using three models: top-down predictive filling-in alone, pRF shift/expansion alone, and a combination of predictive filling-in alone + pRF shift/expansions, and found that including predictive filling-in greatly reduced the magnitude of pRF shifts/expansions. Our results suggest that claims of cortical reorganization based on pRF location or size changes near scotoma boundaries, where predictive filling-in has not been accounted for, should be treated with caution.

## Methods

### Participants

Data were acquired from 10 participants (6 females, 4 males), ages 20 – 38 (*M* = 28.1, *SD* = 6). All participants were paid and had normal or corrected-to-normal acuity. All gave informed consent in accordance with the human participant Institutional Review Board at the University of Washington, in adherence with the Declaration of Helsinki. All 10 participants are a subset of the 12 participants that contributed to a previous study that introduced the log-bar stimulus (Chang et al., 2025). The advantage of this log-bar stimulus is that the conventional drifting bar stimuli used in pRF estimation undersample the central visual field, where receptive fields are small and cortical magnification is high.

### Stimuli

Stimuli were presented on a BOLDscreen32 LCD monitor (Cambridge Research Systems, Rochester, UK) operating at a resolution of 1920 × 1080 at 60 Hz with a viewing distance of 133 cm. An Intel i7 Mac Mini running MATLAB R2022a (MathWorks Inc., Natick, MA) controlled custom stimulus presentation code based on Psychtoolbox-3 (Brainard, 1997; Pelli, 1997). Behavioral responses were recorded using a button box (Current Designs, Haverford, PA, USA).

All stimuli consisted of sweeping bar apertures containing an 8 Hz, 1 cpd contrast-reversing checkerboard. Unlike traditional pRF mapping stimuli that use a bar of uniform width, we used a log-bar stimulus in which the bar is logarithmically warped along the eccentricity dimension so as to more evenly sample visual space in cortical coordinates (Chang et al., 2025). This approach improves the precision of pRF estimates in the foveal representation and provides a more accurate basis for modeling responses near the scotoma boundary.

To create the log-bar stimuli, each frame of a fixed-bar stimulus was distorted along its eccentricity axis, expressed as:

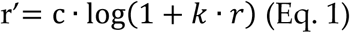

Where *r* is the eccentricity (in degrees of visual angle) of each pixel, k is a distortion factor, *c* is an overall scale constant, and *r’* is the mapped eccentricity of that pixel for the log-bar stimulus. We used a distortion factor of k *=* 5 and a scaling constant of 2.15. This transformation increases sampling density near the fovea, yielding more reliable estimates of pRF position and size, particularly for small receptive fields (Chang et al., 2025). See the OpenNeuro repository for MATLAB/Python code that demonstrates how to warp an image into log-eccentricity space.

Bars were constrained to a central 16o diameter circular aperture. The display was uniform gray outside the aperture. Bars either spread across the entire circular aperture – the ‘full’ stimulus, or had the central 2o replaced by the gray background – the ‘scotoma’ stimulus.

Figure 1. shows example frames from a single sweep of the ‘full’ and ‘scotoma’ stimuli used in these experiments.

**Figure 1.**
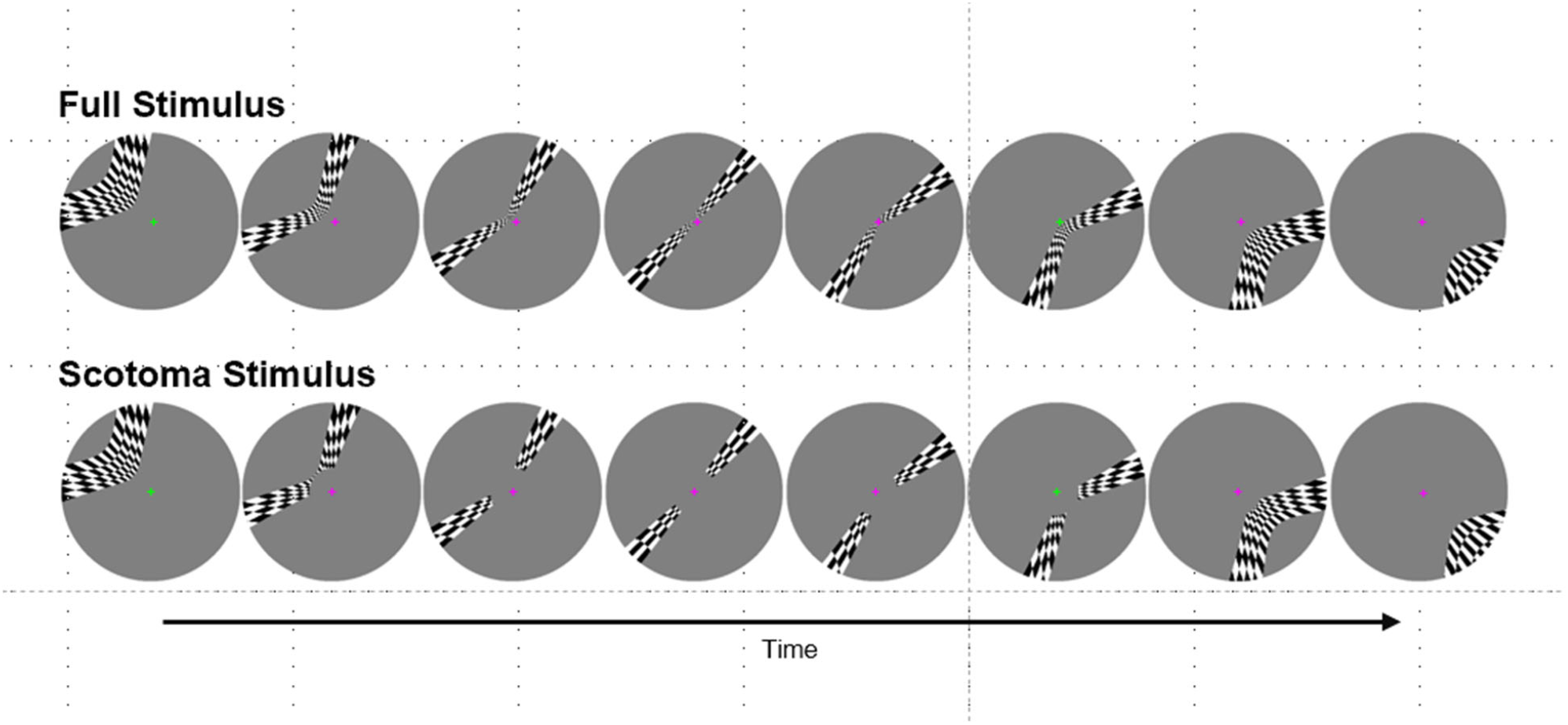
Example frames from the ‘full’ and ‘scotoma’ stimuli which were produced by distorting a fixed-bar stimulus along the eccentricity axis, see Chang et al. (2025) for details. Participants were instructed to fixate on the central cross and report with a button press when the cross changed color.

Each run began with a 2s blank period, followed by eight 45s sweeps of the bar across the full aperture. Each run ended with a 4s blank period, resulting in a total duration of 366s per run.

All participants participated in 2 one-hour scanning sessions, with up to a 2-week interval between sessions. Each session consisted of 9 runs:

Six runs of the full log-bar stimulus were used to identify population receptive fields (a subset of previously published data Chang et al., 2025). The bar sweep direction across each of these runs was based on a pseudorandom seed that was identical across all 10 participants.

Paired sets of 2×3 runs of the full and scotoma stimulus were used to carry out the modeling of the current study. Critically, we used three pseudorandom seeds which were the same across all 10 participants. For each random seed we collected 2 full-field and 2 scotoma runs which were averaged. This resulted in 3 matched paired time-courses for full field and scotoma sequences, in which the only difference between the protocols was the presence of the scotoma.

### Experimental Task

A small cross (“+”) was presented at the center of the display and was present continuously throughout all experimental runs. The color of the cross switched from green to magenta on average of 15 times per run, with a minimum inter-switch delay of 3.2s. Participants were instructed to maintain fixation on the cross and to press a response button whenever the color of the cross changed. The purpose of this task was to encourage fixation for the entire scan duration and focus attention on the fixation spot rather than the moving bar stimulus. Eye-position was tracked in the scanner using an MR-compatible eye-tracking system (SR Research Eyelink 1000 Plus, Ottawa, Canada).

### MRI Acquisition

MRI data were acquired at the University of Washington Center for Human Neuroscience (CHN) on a Siemens 3T Prisma with a 64-channel phase-array head coil. The protocol included 3D volumetric navigator-guided T1-weighted (T1w, multiecho) and T2-weighted (T2w) images acquired at 0.8 × 0.8 × 0.8 mm isotropic resolution (FOV = 166.4 × 240 × 256 mm3, matrix size = 208 × 300 × 320; Tisdall et al., 2012).

BOLD data were acquired using a T2*-weighted 2 × 2 × 2 mm isotropic multiband gradient-echo (multiband acceleration = 4), echo-planar imaging sequences (TR / TE = 1200/30 ms, flip angle = 64o, FOV = 212 × 212, matrix size = 106 × 106, 56 oblique axial slices). Each run collected 305 volumes (366s duration). We collected a total of 9 runs (6 full-field, 3 scotoma) from each subject across two sessions for each participant. Additionally, a pair of opposite phase-encoded EPI references were acquired for each session.

### Preprocessing

Data were processed using the fMRIPrep-21.0.1 pipeline (Esteban et al., 2019). The following details are based on fMRIPrep’s documentation. The T1w image was corrected for intensity non-uniformity with N4BiasFieldCorrection (ANTs-2.3.3; Tustison et al., 2010) and used as the T1w-reference throughout the workflow. The T1w-reference was then skull-stripped with a Nipype implementation of the antsBrainExtraction.sh workflow. Brain tissue segmentation of cerebrospinal fluid (CSF), white matter (WM), and gray matter (GM) was performed on the brain extracted T1w using FSL’s fast segmentation tool (Zhang et al., 2001). Brain surfaces were reconstructed using FreeSurfer recon-all (Dale et al., 1999).

A B0-nonuniformity map (or field map) was estimated based on two echo-planar imaging (EPI) references with topup from FSL (Andersson et al., 2003). A field map was estimated for each session, and the correction was applied to all functional data within the collected session.

BOLD runs were slice-time corrected to 0.549s (50% of the slice acquisition range 0s – 1.1s) using 3dTshift from AFNI (Cox, 1996). The BOLD reference was then co-registered with the T1w reference using bbregister (FreeSurfer), which implements boundary-based registration (Greve & Fischl, 2009). Co-registration was configured as a rigid-body transformation with six degrees of freedom. The BOLD time series was then resampled onto participant-specific surface space in a non-gridded resampling performed using mri_vol2surf (FreeSurfer).

Head-motion parameters with respect to the BOLD acquisitions were estimated and expanded by including temporal derivatives and quadratic terms for each parameter (Satterthwaite et al., 2013), and a set of physiological regressors was extracted to allow for component-based noise correction (CompCor, Behzadi et al., 2007). After high-pass filtering the preprocessed BOLD time series (using a discrete cosine filter with 128s cut-off), principal components were estimated for the anatomical variant (aCompCor). The aCompCor components were calculated separately within the WM and CSF masks.

All BOLD data underwent temporal high-pass filtering (Fourier GLM, cut-off: 3 cycles) and were denoised with 24 head-motion parameters (rigid-body, temporal derivative, quadratic terms (Satterthwaite et al., 2013) and the first 5 aCompCor (WM + CSF) components (Behzadi et al., 2007)). Voxel time-courses were set to have a zero mean.

### Region of Interest (ROI) Selection

Initial visual cortex region selection for pRF fitting was performed by aligning each participant’s anatomical surface with Benson’s atlas of the visual cortex using neuropythy (Benson & Winawer, 2018). The entire visual cortex mask was dilated to ensure the regions of interest (ROI) encompassed the edges of each visual area. This process created a visual cortex region of interest for each participant and hemisphere used in the pRF fitting process.

Benson’s atlas was used again after pRF fitting to fine-tune the visual area boundaries by including participants’ pRF information from all stimulus types and runs. Finally, manual examination and editing were performed to create final V1, V2, and V3, ROIs for each individual.

### Population Receptive Field (pRF) Estimation

PRFs were estimated using methods adapted from Dumoulin and Wandell (2008). The pRF for each voxel, *v*, was modeled as two-dimensional Gaussian function *G*_*v*_ parameterized by a center location *(x, y)* and size, *σ*, (all units in degrees of visual angle). This Gaussian was used to predict this voxel’s BOLD response to the full stimulus using a forward model, which is the overlap between the stimulus and the Gaussian pRF followed by a convolution with a hemodynamic response function (HRF).

Specifically, for a given voxel *v*, and Gaussian pRF *G*_*v*_*(x, y)*. If S*(x, y, t)* is the spatial and temporal aperture for a full field stimulus, then the voxel’s response is predicted by the ‘forward model’ to be:

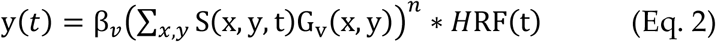

Where * denotes convolution and β_*v*_ represents the scaling of the response amplitude. The center and size parameters, x, y, σ were estimated using a two-stage procedure consisting of a coarse grid search followed by nonlinear optimization to maximize the correlation between predicted and measured time courses (see (Chang et al., 2025) for more detail).

For the linear model, used for the majority of analyses, n was fixed at 1. For the compressive spatial summation model n was an additional free parameter that was allowed to range between 0.05 *<* n *<* 1.35. Multiple starting values were used for n to reduce sensitivity to local minima.

As described above, 6 independent runs of the log-bar full field stimulus was used to select voxels for analysis. Voxels were retained if their pRF centers fell within the stimulus aperture of 8 degrees eccentricity, pRF size exceeded a minimum threshold of 0.1 deg, and the pRF model explained more than 20% of the variance in the measured time series.

### Filling-in model

For each voxel, we denote the measured fMRI responses to the full-field and scotoma stimuli as R_full_(t) and as *R*_scot_*(t)* respectively. Using each voxel’s pRF and the known stimulus sequences, the forward model generated the corresponding predicted bottom-up time-courses *B*_full_*(t)* and as *B*_scot_*(t)*.

The portion of the predicted response to the full field stimulus that is due to the scotoma region can be estimated as Δ*B(t) = B*_full_*(t)* − *B*_scot_*(t)*.

Because the output of the forward model is in arbitrary units, each voxel was assigned a scaling parameter, *β*. This parameter was estimated by scaling the full-field prediction *B*_full_*(t)* to the measured full-field response *R*_full_*(t)*.

We next estimated the fraction, *k*, of the missing response, Δ*B(t)*, that was restored due to top-down feedback that represents the scotoma region. The predicted response the scotoma stimulus was estimated as:

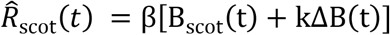

When k=0, the predicted scotoma response is determined entirely by the bottom-up response to the scotoma stimulus:

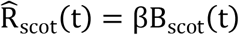

When k =1, the stimulus is completely filled in. Because Δ*B(t) = B*_full_*(t)* − *B*_scot_*(t)* it follows that:

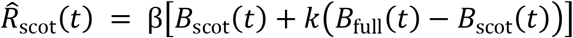

Which reduces to:

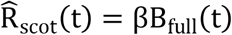

Thus, the predicted response to the scotoma stimulus is equal to bottom-up response to the full stimulus:

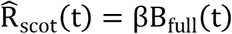

Intermediate values of *k* indicate partial restoration of the response that would otherwise be lost because of the scotoma.

A single value of *k* was estimated for each participant, visual area, and 0.25° eccentricity bin. Within each bin, the voxel-specific scaling parameters *β*_*v*_ were fixed at the values estimated from the full-field data, while *k* was shared across voxels. Bounded least-squares minimization, with 0 ≤ *k* ≤ 1 was used to find the value of *k* that minimized the summed squared error between the predicted and measured scotoma responses across all retained voxels and time points. This procedure yielded one estimate of *k* for each participant, ROI, and eccentricity bin.

## Results

### Behavioral Task

Participants’ responses to the color detection task were analyzed across 12 functional runs (including three runs not included in this paper). Performance was defined as:

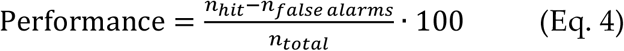

where *n*_*hit*_ is the number of successful color change detections (defined as a button press within 1s of a color change), *n*_*false alarms*_ indicates the number of additional responses without a color change, and *n*_*total*_ indicates the total number of color changes. Average behavioral performance was 96.39%, and the interquartile range of performance values across participants was (95.28%, 98.89%).

### Filling-in

As described in Methods, the parameter, *k*, represents the proportion of the ‘missing’ response in a scotoma that is replaced by top-down filling-in. **Figure 2A** shows k as a function of eccentricity for voxels in areas V1, V2 and V3. Because the parameter k represents the proportion of the *missing* signal that is filled in, it rapidly becomes undefined outside the scotoma where there is no ‘missing’ signal to be filled in.

**Figure 2.**
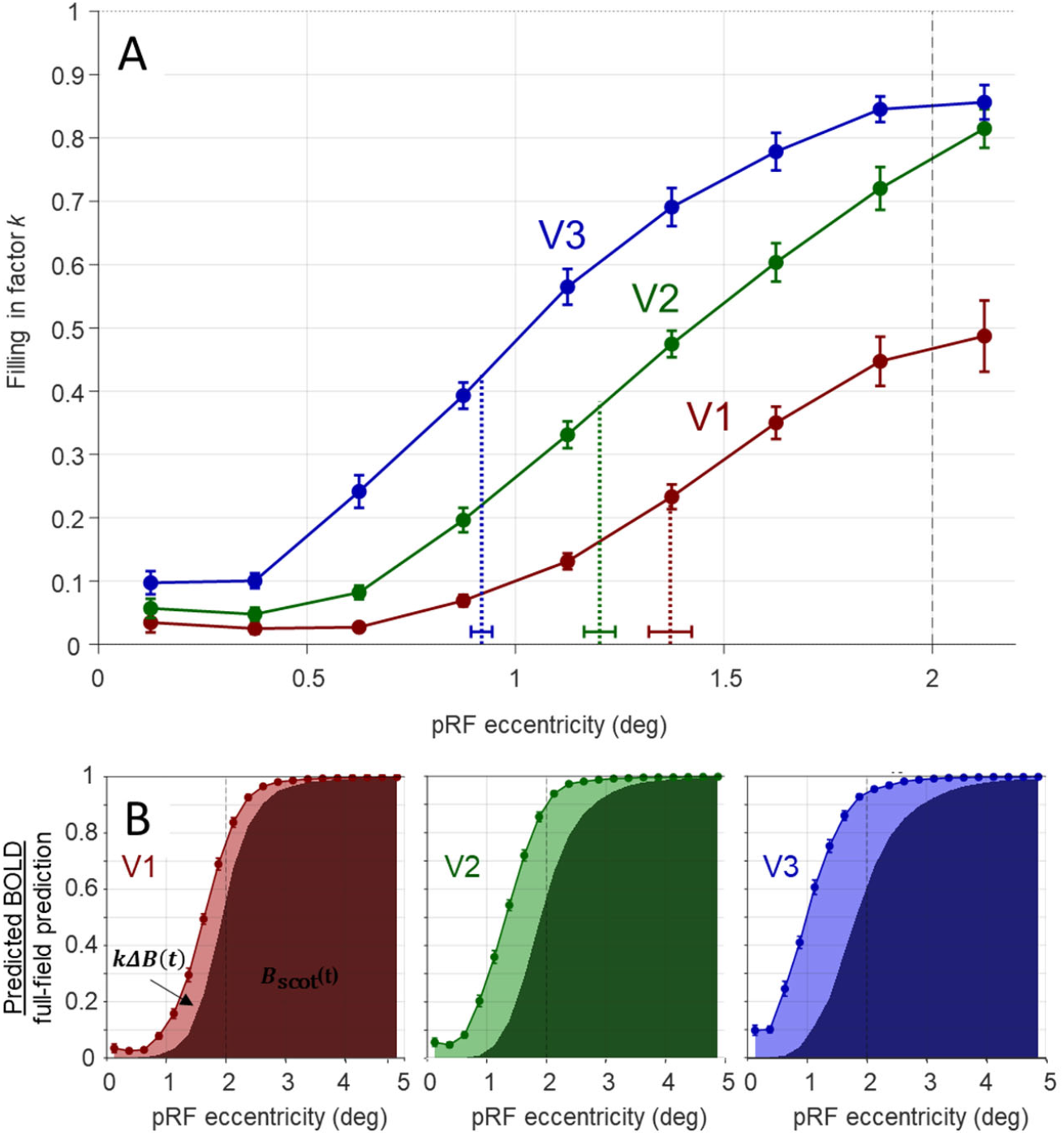
(A) Filling-in factor k as a function of eccentricity in areas V1 (red), V2 (green) and V3 (blue). Symbols are means, error bars are SEM, and small points are individual subjects. Gray region indicates eccentricities outside the artificial scotoma. Half-width at half-height was calculated using linear interpolation, with k at 2 degrees as the reference. (B). The relative contributions of spilling-in (dark shading, s) and top-down filling-in (light shading, f = k × (1-s)) for areas V1-V3.

Three interesting results can be observed, which are supported by fitting a linear mixed-effects model with fixed effects of eccentricity, ROI, and their interaction, and a random intercept for subject.

First, filling-in strength increased reliably with eccentricity within all three visual areas: (V1: F(7,72) = 101.67, p<.0001, V2, (F(7,72) = 231.04, p<.0001), V3 (F(7,72) = 256.87, p<.0001). In V3, *k* reaches values above 0.8 near the edge of the scotoma, suggesting that almost all the missing signal is filled in.

Second, near the fovea (eccentricities below 0.75), *k*-values are significantly greater than zero in each visual area (V1: mean k = 0.029, t(9) = 3.50, p = 0.0067; V2: mean k = 0.062, t(9) = 5.99, p = 0.0002; V3: mean k = 0.146, t(9) = 9.18, p < .0001) indicating that there is some filling-in even for voxels with pRFs well inside the artificial scotoma.

Finally, *k*-values increase across the visual hierarchy from V1 to V3. This is seen as a robust interaction (k ~ ecc*ROI + (1|subject)) between eccentricity and ROI (F(14,216) = 22.14, p < .0001), indicating that the eccentricity profile of filling-in differs across areas. We also found that the extent of filling-in increased across the hierarchy, as shown by the half width at half height values for V1-V3 (V1 = 0.63 +/− 0.05 deg, V2 = 0.80 +/− 0.04 deg, V3 = 1.08 +/− 0.03 deg).

Another useful way of formulating predictive filling-in is to ask what proportion of the neural response is due to top-down filling in? Panel B shows stacked curves, with the lower curve showing the proportion of the neural response driven by bottom-up spilling-in, *B*_scot_(t). This spilling-in response is driven by pRFs that at least partially lie outside the scotoma.

The upper curve represents the additional response driven by top-down filling-in: *k*Δ*B(t)*. Differences between the upper curve and y *=* 1 represent the drop in neural signal within the scotoma.

### Estimates of filling are robust to model parameters

Using our linear model, we observe robust filling-in across all three visual areas. How robust are our estimates of filling-in to errors or variations in our pRF model?

#### Errors in pRF size

First, we determined the robustness of our filling-in estimates to potential misestimation of pRF size. The mid-shaded lines in Figure 3 show a re-estimation of the filling-in parameter *k* after doubling pRF sizes (*σ* × 2) while keeping the pRF center location fixed. These larger pRFs were then used in the forward model to re-generate predicted responses to the full and scotoma stimuli. As expected, larger pRFs increased estimates of spilling-in, but estimates of *k* were surprisingly robust to misestimations of pRF size.

**Figure 3.**
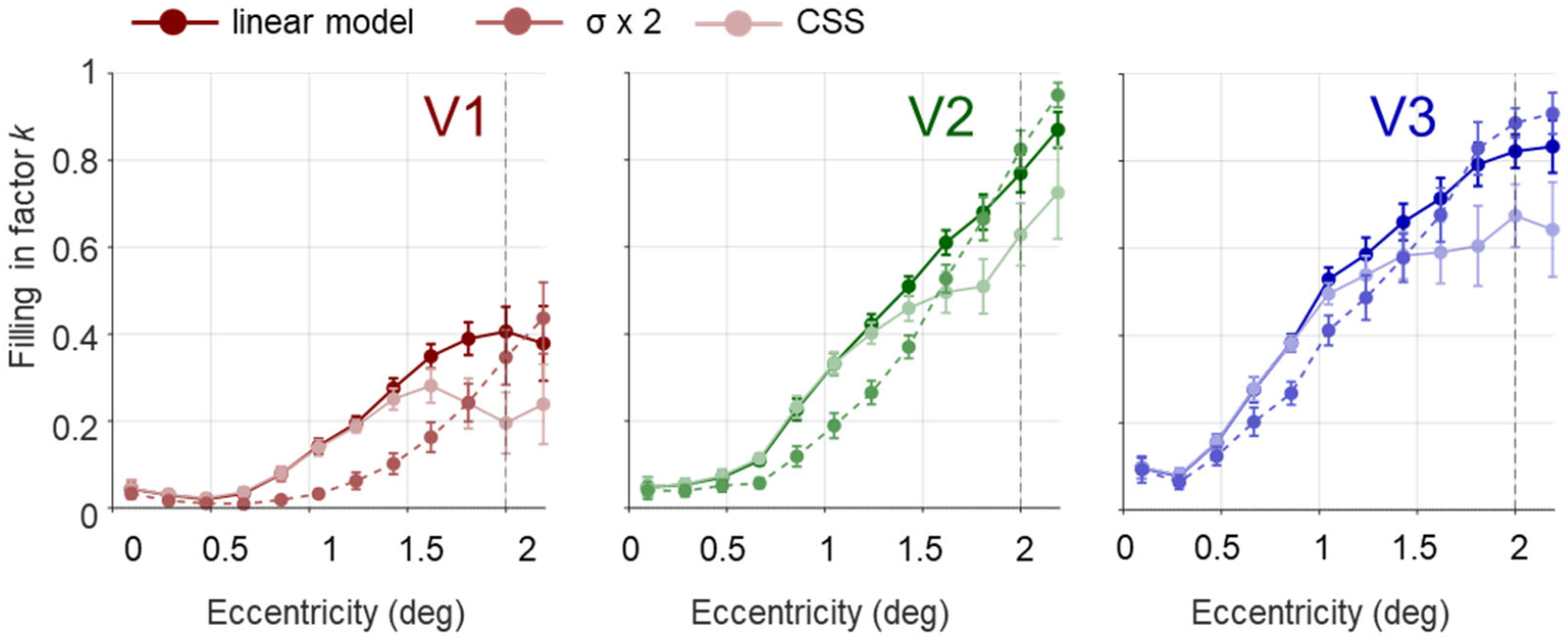
Filling-in factor k as a function of eccentricity in areas V1 (red), V2 (green) and V3 (blue). Dark curves represent the original linear pRF model. Mid-shaded curves represent estimates made using a forward model with receptive field sizes ofσx 2. Light curves represent estimates of k using the compressive spatial summation model.

#### Compressive spatial summation

Next, we examined filling-in using a nonlinear pRF model in which n was allowed to vary, in a model previously described by (Kay et al., 2013). The lightest lines in Figure 3 show estimates from this model - we see significant filling-in. For the CSS model we see a sharp decline in k near the scotoma border. This is likely because the compressive nonlinearity was also applied to the filling-in component. As a result even a small amount of top-down filling-in signal results in a large increase in the predicted BOLD response.

### Predictive filling-in vs. changes pRF location and size

A variety of studies have reported substantial shifts in pRF location or size near the boundaries of scotoma (Binda et al.; Carvalho et al.; Hummer et al., 2018). Critically, these shifts are observed using predictable stimuli and forward models that did not include the effects of top-down filling-in.

We fit our data using three models: our linear top-down predictive filling-in model, a model where pRFs were allowed to shift/expand and we assumed no filling-in (setting k=0) - similar to (Binda et al., 2013; Carvalho et al., 2021; Hummer et al., 2018), and a model that included both predictive filling in and shifts/expansions of receptive fields.

The upper panel of Figure 4 compares shifts in pRF locations across the model where pRFs were allowed to shift/expand and we assumed no filling-in and the model that included both predictive filling in and shifts/expansions of receptive fields. While both models show shifts in receptive fields, the shifts are much larger for the model that does not incorporate filling-in, suggesting that some of the shifts in location observed in previous studies might reflect top-down filling in rather than changes in pRF profiles. Similarly, changes in size (middle panel) are much smaller when predictive top-down effects are taken into account. Thus, claims of cortical reorganization based on pRF location or size changes near scotoma boundaries, where predictive filling-in has not been accounted for, should be treated with caution.

**Figure 4.**
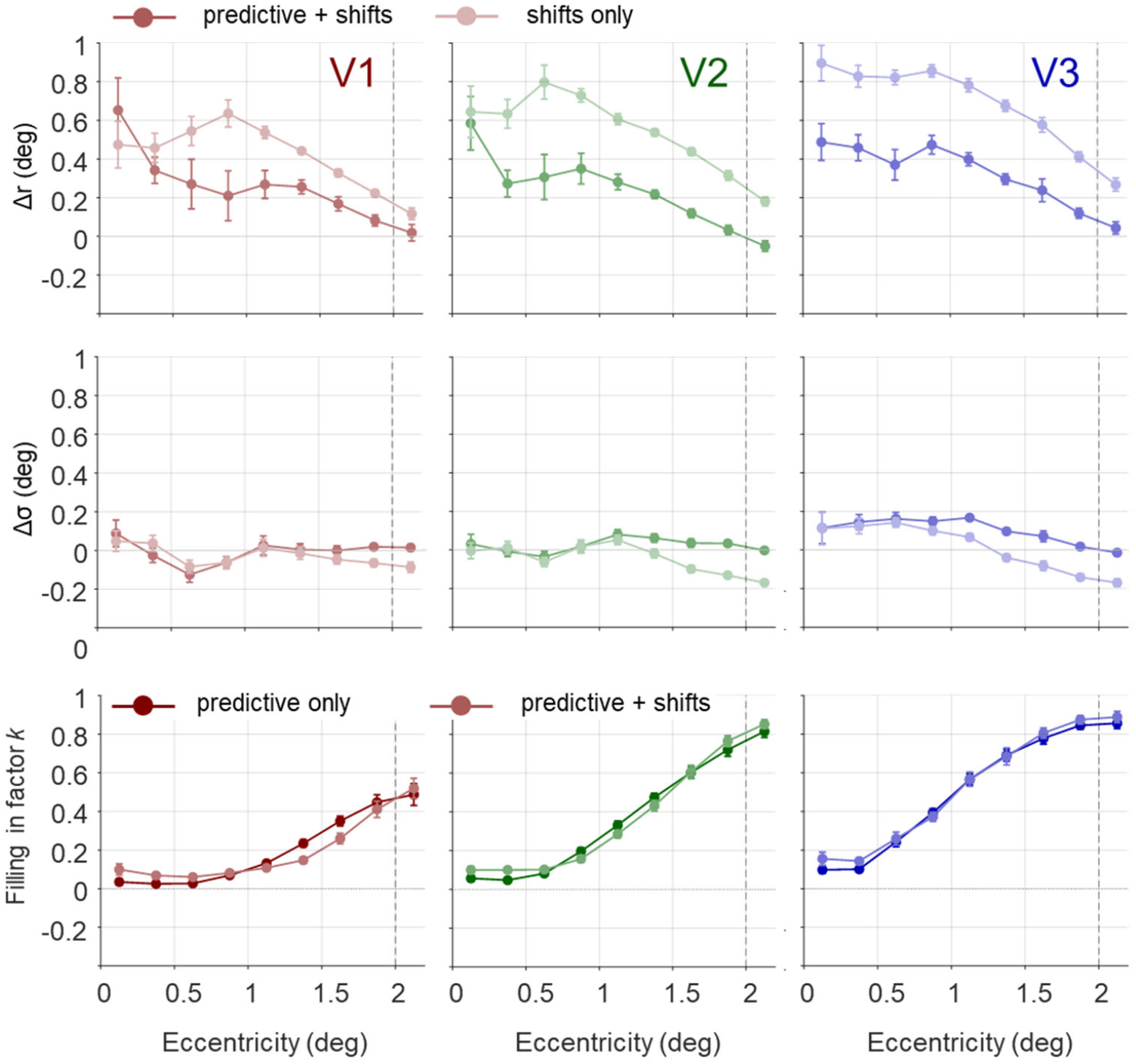
Filling-in factor k as a function of eccentricity in areas V1 (red), V2 (green) and V3 (blue). Dark curves represent the original linear pRF model. Mid-shaded curves represent estimates of k using a forward model with receptive field sizes ofσx 2. Light curves represent estimates of k using the compressive spatial summation model. Note that estimates for eccentricities smaller are unreliable, because the signal is too small to accurately estimate pRFs. Upper Panels: Shifts in pRF locations for full vs. scotoma condition. Positive values represent a shift outward. Middle Panels: Shifts in pRF size for fill vs. scotoma conditions. Lower panels: Estimates of k when pRFs are fixed vs. allowed to shift.

Finally, the bottom panel compares estimates of *k* in our original model to estimates based on a model that includes pRF shifts. Estimates of *k* are similar – failing to model top-down filling-in responses can result in spurious shifts in pRF location, but failing to model changes in pRF location does not seem to significantly impact estimates of predictive filling-in.

## Discussion

Many previous studies have observed robust BOLD responses within voxels whose receptive field centers lie within a scotoma, and as described in the introduction, these responses within a scotoma may be caused by any or all of the following: stimulus-driven responses outside the scotoma that are carried into the region – ‘spilling in’, they may reflect top-down filling-in (Meng et al., 2005; Muckli et al., 2015; Papale et al., 2023; Smith & Muckli, 2010; Tong & Engel, 2001; Weerd et al., 1995) or they may reflect cortical reorganization in which pRF locations shift or expand to include regions outside the scotoma (Barton & Brewer, 2015; Baseler et al., 2011; Binda et al., 2013; Carvalho et al., 2021; Dumoulin & Knapen, 2018; Ferreira et al., 2017; Gilbert & Wiesel, 1992; Haak et al., 2012; Hummer et al., 2018; Kaas et al., 1990; Meng et al., 2005; Papanikolaou et al., 2015; Prabhakaran et al., 2020). The present study introduces a model-based approach for separately quantifying these three factors.

The first important finding of our study is that responses from well within a scotoma can arise from stimulation outside its boundaries. As shown in Figure 2, a substantial portion of the BOLD response of voxels within a scotoma can be explained by pRF overlap alone.

However, even after accounting for this effect, a residual “filling-in” signal remains which increases systematically across the visual hierarchy, from V1 to V2 to V3. This pattern is consistent with prior work suggesting that completion-related signals become more prominent in extrastriate cortex (Mendola et al., 1999; Weerd et al., 1995), but our study is the first to quantify differences in filling-in across the visual hierarchy. This finding that filling-in increases across the hierarchy is consistent with predictive coding frameworks, in which higher visual areas generate predictions about sensory input that are fed back to earlier areas (Friston, 2005; Rao & Ballard, 1999).

Finally, we show that failing to incorporate filling in is likely to result in an overestimation of cortical reorganization. While we did see some evidence of systematic shifts in pRF locations towards the edge of the scotoma, these shifts were approximately half the size of the shifts seen when predictive signals were not included in the model.

Why should failing to account for top-down filling-in produce an apparent shift in pRF position? Consider a voxel whose receptive field straddles the scotoma border. The portion of the pRF inside the scotoma receives no bottom-up input during presentation of the scotoma stimulus, but top-down input results in the voxel responding similarly to when the bar has swept through it. The only way a forward model with a Gaussian pRF can account for this extra signal is to displace the pRF center away from the scotoma or by enlarging it. For many voxels near the scotoma border there is likely to be some new position and size that produces a response over time that more closely matches a neural response that includes filling-in. Because displacement parameters can absorb the filling-in signal while producing good fit values, failing to include filling-in is likely to result in spurious overestimates of cortical reorganization.

Interestingly, allowing pRFs to shift and expand barely altered our estimates of k. This is because filling-in enters the model as a single fixed regressor that cannot easily be accounted with shifts in pRF location.

Two limitations of the present study should be noted. First, our pRF model assumes a simple Gaussian receptive field and a linear (or simple nonlinear) relationship between stimulus and response. While this model captures the coarse spatial properties of voxel responses, it does not account for nonlinearities such as surround suppression or gain modulation, which may influence responses near the scotoma boundary. Second, our estimates of filling-in depend on the accuracy of pRF fits and the forward model predictions; systematic errors in these estimates could bias inferred values of *k*. However, it should be noted that our general pattern of results seems to be robust to our estimates of pRF size and the exact forward model architecture.

The ability to dissociate bottom-up spilling-in from top-down filling-in and shifts in pRF locations and size has important implications for interpreting cortical responses in clinical populations with visual field defects. For example, training-induced improvements in detection performance have been linked to spared activity in regions of V1 adjacent to the lesion (Barbot et al., 2021). In these patients recovered visual function is often associated with activity in perilesional visual cortex – exactly the location where top-down filling in has the strongest influence on the neural response and can produce the strongest effect of spurious changes in pRF location. Our data suggest that interpretating this activity correctly requires disentangling the degree to which responses in perilesional cortex reflect an upweighting of residual input from intact regions of the visual field, enhanced top-down filling-in, and changes in pRF location and size.

## Acknowledgments

This research was supported by grants from NIH National Eye Institute Grant R01EY014645 (IF) and the University of Washington Center for Human Neuroscience (CHN).

The log-bar warping method reported in this article is available in the OpenNeuro repository (https://doi.org/10.18112/openneuro.ds004698.v2.0.0).

